# Nitric oxide-dependent inflammation underlies Notch and PI3K/Akt oncogene cooperation

**DOI:** 10.1101/174169

**Authors:** Santiago Nahuel Villegas, Rita Gombos, Irene Gutiérrez Pérez, Lucia García López, Jesús García-Castillo, Diana Marcela Vallejos, Vanina Da Ros, József Mihály, Maria Dominguez

**Affiliations:** Instituto de Neurociencias, Consejo Superior de Investigaciones Científicas (CSIC) and Universidad Miguel Hernandez (UMH), Campus de Sant Joan, Apartado 18, 03550 Sant Joan, Alicante, Spain; Institute of Genetics, Biological Research Centre HAS, MTA-SZBK NAP B Axon Growth and Regeneration Group, Szeged, Hungary

**Keywords:** chemical screen, Cancer, Drosophila, Notch, PI3K/Akt, Pten, inflammation, nitric oxide synthase, lipoxygenases

## Abstract

Concurrent activating mutations of the Notch and PI3K/Akt signalling pathways cooperate in the induction of aggressive cancers. Unfortunately, direct targeting of any of these aberrant pathways can result in severe side effects due to their broad physiological roles in multiple organs. Here, using an unbiased chemical *in vivo* screen in *Drosophila* we identified compounds that suppress the activity of the pro-inflammatory enzymes, nitric oxide synthase (NOS) and lipoxygenase (LOX), capable to block oncogenic Notch-PI3K/Akt cooperation without unwanted side effects. Genetic inactivation of NOS and LOX signalling components mirrors the anti-tumorigenic effect of the hit compounds. We show that NOS activity and immunosuppression associated to inflammation facilitates Notch-mediated tumorigenesis. Our study reveals an unnoticed immune inflammatory process underlying Notch-PI3K/Akt tumours and exposes NOS as a druggable target for anti-cancer therapeutic development.

## Introduction

Crosstalk between activated Notch and the phosphatidylinositol 3-kinase (PI3K)/Pten/Akt pathways increases proliferation, survival and invasion of cancer cells^1-3^. This oncogenic combination is highly aggressive and is found in both liquid^2,4,5^, and solid cancers^1,6,7^. Combination of Notch and Akt inhibitors effectively kills cancer cells and bypasses individual pathway inhibition resistance. However, as these signalling pathways traverse numerous biological functions their pharmacological systemic inhibition seriously hamper whole organism physiology^8,9^, resulting in severe short- and long-lasting side effects^10-12^. These observations highlight the necessity for exploring safer alternatives and therapeutic approaches that indirectly targets this oncogene cooperation.

*Drosophila* is a proven model system for cancer research^13-18^ and provide a potent *in vivo* system for phenotype-based drug screening^19-22^ that can be extrapolated into mammalian pharmacology^13^. Moreover, the fact that both the Notch and the PI3K/Akt signalling cascades are highly conserved from flies to humans^23-25^, together with the well-studied role of these individual pathways in fly tumourigenesis^26-29^ makes *Drosophila* a particularly useful model for anticancer drug testing aiming to target this oncogenic cooperation.

In this study, we used a *Drosophila* eye cancer model of Notch and PI3K/Akt cooperative oncogenesis to perform an *in vivo* drug screen with the aim to identify compounds capable to suppress tumorigenesis without causing side effects to the treated animals. We found that the anti-inflammatory drug BW B70C and related compounds with the capacity to inhibit the nitric oxide synthase (NOS) and LOX pathways, blocks Notch-PI3K/Akt induced tumorigenesis. Genetic analysis further defined the role of NOS/LOX inflammatory signalling pathway in tumour growth induced by the Notch-PI3K/Akt cooperative action. We show that Akt-dependent activation of NOS or mimicking an inflammatory pro-tumorigenic environment by blocking the immune system response is sufficient to unleash Notch oncogenic potential. Collectively, our study unveils NOS as a potential safe pharmacological target and places a class of anti-inflammatories as promising compounds for the development of therapeutic approaches to block Notch-PI3K/Akt cooperative oncogenic action.

## Results

### Unbiased chemogenomic in vivo screen for targeting Notch-PI3K/Akt oncogenic cooperation

We employed an unbiased, phenotype-based chemical screen with the goal of identifying new agents that blocks Notch-PI3K/Akt oncogenic cooperation without harming normal cells. Two *Drosophila* eye cancer models that capture the molecular features of Notch-PI3K/Akt cooperative oncogenesis were used (Fig. 1a and Supplementary Fig. 1a). Activation of Notch signalling was induced by the overexpression of its ligand *Delta* (*Dl*) and to hyper-activate the PI3K/Akt pathway we used either overexpression of Akt^2,17^ or interference RNA (RNAi) to silence Pten, an Akt negative regulator. The eyeless (ey) promoter was used to drive concurrent eye-specific overexpression during a tightly regulated developmental period in the eye imaginal disc. The cooperative action of these pathways is what causes the development of eye tumours, as the sole activation of either the Notch (*ey>Dl*) or PI3K/Akt (*ey>Akt*) pathways is not sufficient to promote tumorigenesis^2,17^ (Fig. 1a). The *ey>Dl>Akt* and *ey>Dl>Pten-RNAi* models yield a similar and robust eye tumour phenotype (Fig. 1a and Supplementary Data 1a) that is readily observed in live anaesthetized adult flies. As the tumour incidence is about 70% (Supplementary Fig. 1a) the model allows the identification of compounds that either suppress or enhance the tumorigenic phenotype. We first observed that systemic inhibition of Notch using the γ-secretase inhibitor DAPT suppresses tumorigenesis but also interferes with physiological cell and tissue growth, consequently resulting in ‘notched’ wings and high lethality (Fig. 1a and Supplementary Fig. 1b). Similarly, systemic inhibition of PI3K/Akt signalling using the PI3K inhibitor LY294002 or combined concurrent block of the two pathways resulted in severe lethality (Supplementary Fig. 1b), thus, indicating comparable toxic side effects as seeing in human cells^7^.

**Figure 1.**
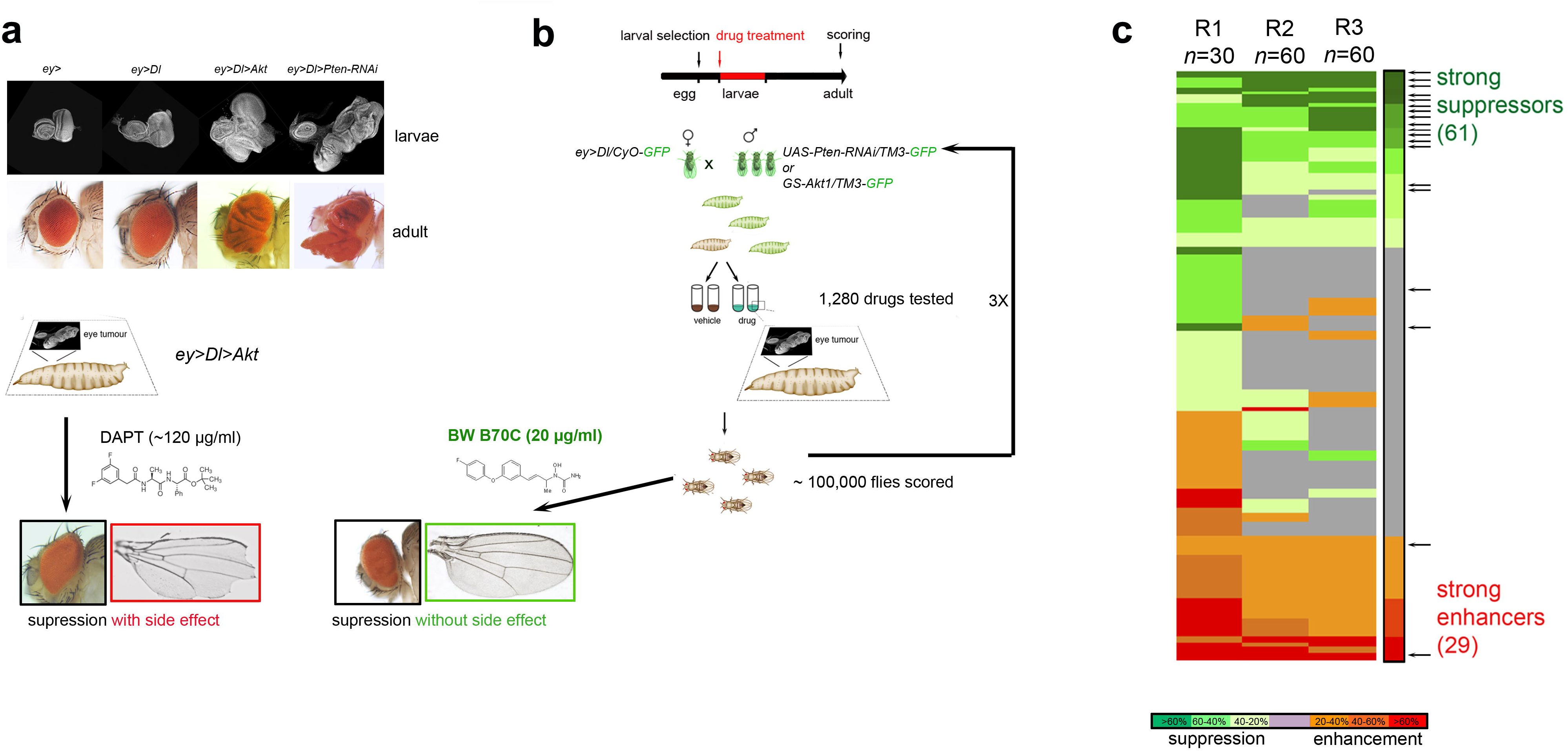
*In vivo* screen identified compounds targeting Notch-PI3K/Akt cooperative oncogenesis. (**a**) Larval eye imaginal discs (top) and adult eyes (bottom) of wild type and tumorigenic flies. Tumorigenesis was induced synergistically through the co-overexpression of *Dl* with either *Akt-* or *Pten-RNAi* (*UAS-Pten-RNAi*^*BL25967*^) in the proliferating larval eye tissue using *ey-Gal4* (*ey>*). These eye cancer models are suitable for detecting the drug response, and side effects that result from normal Notch pathway inhibition (i.e., ‘notched’ wings). Lower image shows an example of the progeny of γ-secretase inhibitor (DAPT)-treated tumour-bearing larva. (**b**) Schematic diagram of the screen design. Tumour-bearing larvae (non-GFP) were either treated with compounds (100 μg/ml in the food) or with the vehicle control. Lower image shows an example of the progeny of *ey>Dl>Akt* larva treated with 20 μg/ml BW B70C, one of the top hit compounds. (**c**) Heat map of the screen results (right column, mean effect). Green denotes suppression; red, enhancement; and grey, no significant change. Arrows point to known anticancer drugs in the LOPAC^1280^. *n*, number of larvae per drug per round (R).

We screened the Library of Pharmacologically Active Compounds (LOPAC^1280^), comprised of 1,280 small molecules including a set of FDA-approved anticancer drugs that acted as internal controls. Furthermore, an annotated list of the known targets of the LOPAC^1280^ drugs is available enabling the readily transformation of phenotypic screening results into a target-based drug discovery approach^23,30^. We tested each LOPAC^1280^ drug, administered with the food, during the larval period at a concentration of 100 μg/ml in three double-blinded rounds, and subsequently assessed the impact on tumour burden and on normal tissue growth in the adult flies (Fig. 1b). This allowed us to directly evaluate both drug response and side effects. Response was calculated by monitoring non-tumorous eyes after treatment and normalized to vehicle control group reared in parallel (Supplementary Fig. 1c). Compounds that showed a lethal effect during the first screen were re-tested at lower concentrations (20 μg/ml).

After the first round of screening (R1, n = 30 larvae per drug), any compound causing greater than 20% response (198 suppressor and 276 enhancer compounds) (Supplementary fig. 1d) were re-screened using a larger number of animals (n = 60 larvae per drug per round). This significantly reduced the number of false positives and led to a high reproducibility (>80%) between rounds R2 and R3 (Fig. 1c). In such a way, after screening approximately 100,000 tumour-bearing flies, we found 90 hit compounds (Fig. 1c) that strongly (>60% response) suppressed (61 hits) or enhanced (29 hits) tumorigenesis (Supplementary Tables 1 and 2). Representative images in Fig. 1a and 1b show a comparison between the response/side effects of Notch inhibitor DAPT and one of the top hit compounds, BW B70C.

Our screen identified 15 of the 21 known anticancer compounds included in the LOPAC^1280^ library (arrows in Fig 1c and Supplementary Table 3) as suppressors. Out of these, we found 13 to strongly suppress, and two to moderately suppress tumorigenesis. Of the remaining six compounds, we identified two as strong enhancers, two as lethal and two with no effect. The fact that we were able to single out these anticancer drugs, some of which are approved by the FDA for the treatment of leukaemia and solid cancers, confirms the validity of our screen. Apart from acting as an internal control for the screen design, these results also show a strong positive correlation with the response observed in human cancer cells.

### RNAi-based functional validation of drug screen results

The remaining 48 strong suppressors (excluding the 13 with known anti-cancer activity) are previously unappreciated modulators of Notch-PI3K/Akt-driven tumorigenesis. As most of these compounds have a known human molecular target, we validated whether their tissue specific genetic down-regulation can mimic the action of the corresponding hit compounds. We used RNAi transgenic expression^31,32^ (Supplementary Fig. 2a) to target 77 *Drosophila* homologous genes of annotated and predicted hit molecular targets in the context of oncogenic Notch-PI3K/Akt (Supplementary Table 4). We reasoned that an anti-neoplastic effect will cause not only a reduction of the tumour burden but will also increase the survival to adulthood. PI3K-RNAi was used as a blind positive control and tumourigenesis was assessed in adults. As a result, RNAi-based validation of hit molecular targets revealed that anti-tumour activity of most of the compounds was consistent with the drugs acting through conserved biological targets rather than target-indirect effects (Supplementary Fig. 2b,c). This supports that, despite the evolutionary distance from humans, our *Drosophila*-based strategy may be capable of identifying drugs and their clinically relevant targets as highlighted in other previous drug screens of human neurodegenerative and neoplastic diseases^20,21,32-36^.

### PI3K/Akt fuels Notch-driven tumorigenesis through NOS

A survey of the hits classified as strong to medium suppressors in our screen revealed the presence of several anti-inflammatory agents targeting the NOS and the lipoxygenase (LOX) signalling pathways (Supplementary Fig. 3a and Supplementary Table 1). Among the top hits the anti-inflammatory compound BW B70C^37^ drew considerable attention given that showed an strong anti-tumorigenic response at a very low concentration (20 μg/ml) (Figs. 1b and Supplementary Table 1).

We first investigated how NOS signalling contributes to Notch-PI3K/Akt-induced tumorigenesis. Using the *Nos* sensor *Nos*^*MI09718 38*^ we observed aberrant expression of *Nos* within the tumour eye tissue (Supplementary Fig. 3a), an action induced by activated Akt (Supplementary Fig. 3b). Then, we treated *ey>Dl>Pten-RNAi* larvae with L-NAME (NG-nitro-L-arginine methyl ester), a selective NOS inhibitor with documented activity in *Drosophila*^39^ and found a significant suppression of tumour incidence (Fig. 2a). In the same line, genetic downregulation of the single *Drosophila Nos* gene through selective knockdown (*ey>Dl>Pten-RNAi>Nos-RNAi*) or endogenous mutation (*ey>Dl>Pten-RNAi; Nos*^*MI09718/+*^) both suppressed tumorigenesis (Fig. 2a), demonstrating a critical contribution of *Nos* to Notch-PI3K/Akt-driven tumorigenesis.

**Figure 2.**
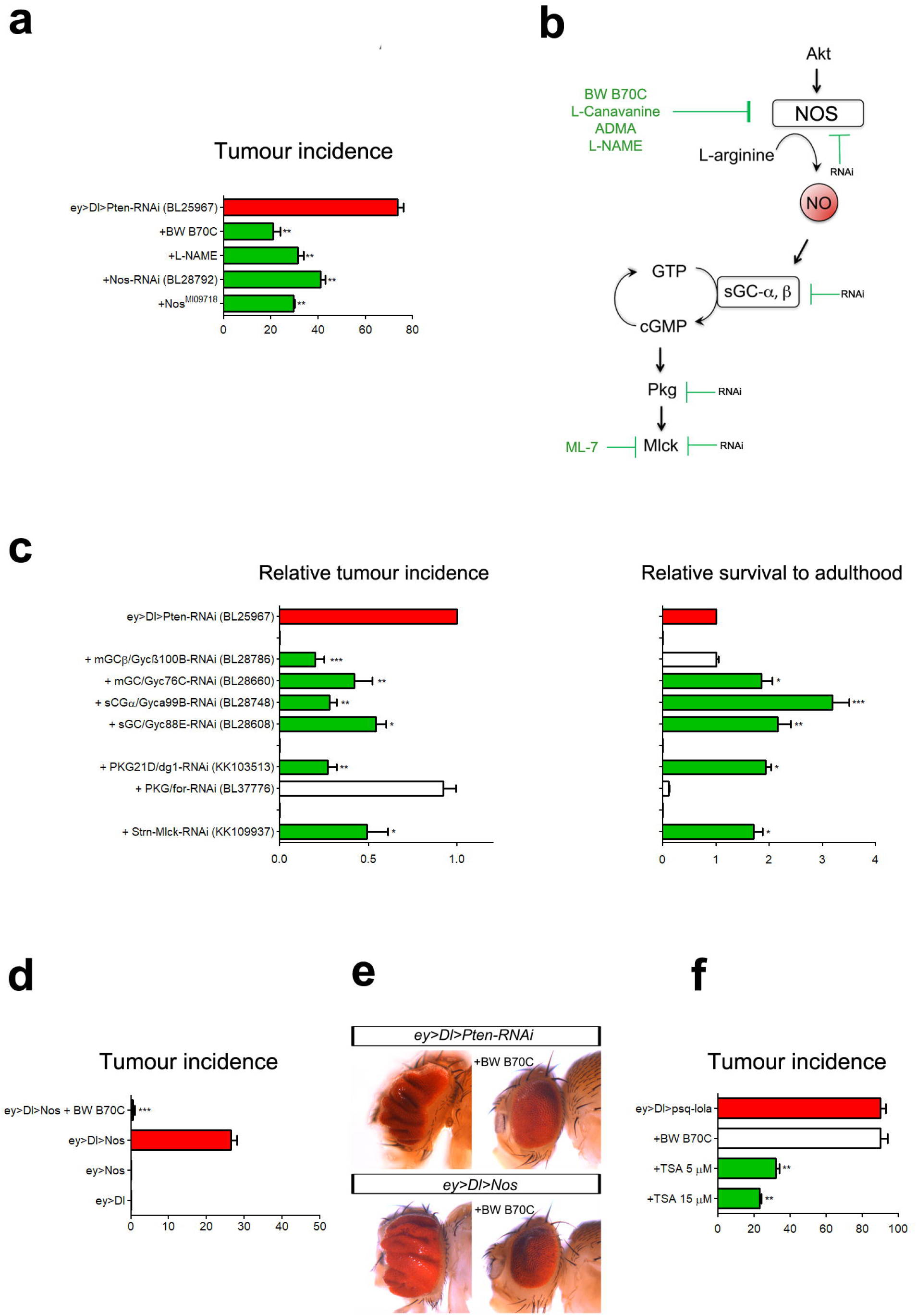
Nos facilitates Notch induced tumorigenesis. (**a**) Graph shows tumour incidence (in %) in flies after pharmacological or genetic inhibition or activation of the indicated genes. (**b**) Schematic NOS signalling pathway. Anti-tumorigenic drugs identified in our screen and RNAi-based silenced genes are shown. (**c**) Effect of RNAi-based silencing of NOS signalling components on tumorigenesis (left graph) and survival (right graph) (*n* = 50-100 eyes per genotype) relative to *ey>Dl>Pten-RNAi*. (d) Tumour incidence (in %) generated by Notch-Nos overexpression (e) Representative *ey>Dl>Pten-RNAi* and *ey>Dl>Nos* eyes treated or not with BW B70C. (f) Tumour incidence (in %) in eyeful tumorigenic model treated with either BW B70C or TSA. \**P<*0.05, \*\**P<*0.01, \*\*\**P<*0.001; One-way ANOVA followed by Bonferroni’s Multiple Comparison test.

Canonical NOS signalling acts through the soluble guanylyl cyclase (sGC)-cGMP-PKG pathway (Fukumura et al, 2006) (Fig. 2b). We found that RNAi-mediated silencing of *Drosophila* genes encoding for the *sGC-α* and *–β* subunits, and *Pkg21D* within tumour cells all suppressed tumorigenesis (Fig. 2c). Furthermore, RNAi silencing of myosin light chain kinase (*Mlck*), a target of PKG, also showed reduced tumorigenesis. This result further validate another of the top hit compounds that we have identified in our screen, ML-7, a potent inhibitor of MLCK (Fig. 2c). In sum, these data imply a tumour cell requirement of *Nos* and highlight the importance of a NO-sGC/cGMP/PKG pathway in Notch-PI3K/Akt-driven tumorigenesis.

Remarkably, overexpression of *Nos* was sufficient to facilitate tumorigenic conversion by Notch independently of PI3K/Akt activation (*ey>Dl>Nos*, Fig. 2d,e). Conversely, eye-specific silencing or overexpression of the *Nos* gene is inconsequential for imaginal eye disc growth^40,41^ and data not shown). This data indicates that Nos activity is sufficient to fuel Notch’s oncogenicity. Importantly, BW B70C was able to block the formation of Notch-Nos-driven tumours (Fig. 2d,e), consistent with the observation that BW B70C treatment decreased Akt-induced *Nos* expression in larvae (Supplementary Fig. 3c). Conversely, tumours induced by the cooperation of Notch with the epigenetic regulators Psq/Lola (the eyeful cancer model)^17^ were not at all sensitive to BW B70C, even though they could be repressed using the epigenetic drug Trichostatin A (TSA; Fig. 2f). Thus, BW B70C does not suppress Notch-driven tumorigenesis in a general manner, but rather seems to inhibit tumour formation by dampening a tumour promoting process orchestrated by Nos aberrant expression.

### LOX pathway inhibition blocks Notch-PI3K/Akt-driven tumorigenesis

LOX enzymatic activity and derivatives have been detected in *Drosophila* extracts and other insects^42,43^, however, the LOX gene(s) still remain undefined. Thus, we searched for *Drosophila* LOX pathway homologous genes that are potentially suitable to validate our screen results further.

Leukotriene A4 hydrolase (LTA4H) catalyses the production of leukotriene B4 (LTB4), a major product of LOX enzymatic activity highly expressed in some cancers^44^. We found that *Drosophila* has a homologous of LTA4H, encoded by *CG10602* (Fig. 3a). Halving the gene dosage of *dLTA4H/CG10602* (*ey>Dl>Pten-RNAi>CG10602*^*f04195*^/+) markedly suppressed tumorigenesis and rescued tumour-associated lethality (Fig. 3b). As leukotrienes exert their action through GPCRs^45^, we also silenced type-A and C allatostatin receptors, the homologous of leukotriene receptors Ltb4r and Cysltr1,2 in *Drosophila* (Fig. 3a and Supplementary Table 4). Inactivation of *AstA-R1* suppressed tumorigenesis, whereas silencing the *AstA-R2*, *AstC-R1*, and *AstC-R2* did not result in tumour growth attenuation (Fig. 3b).

**Figure 3.**
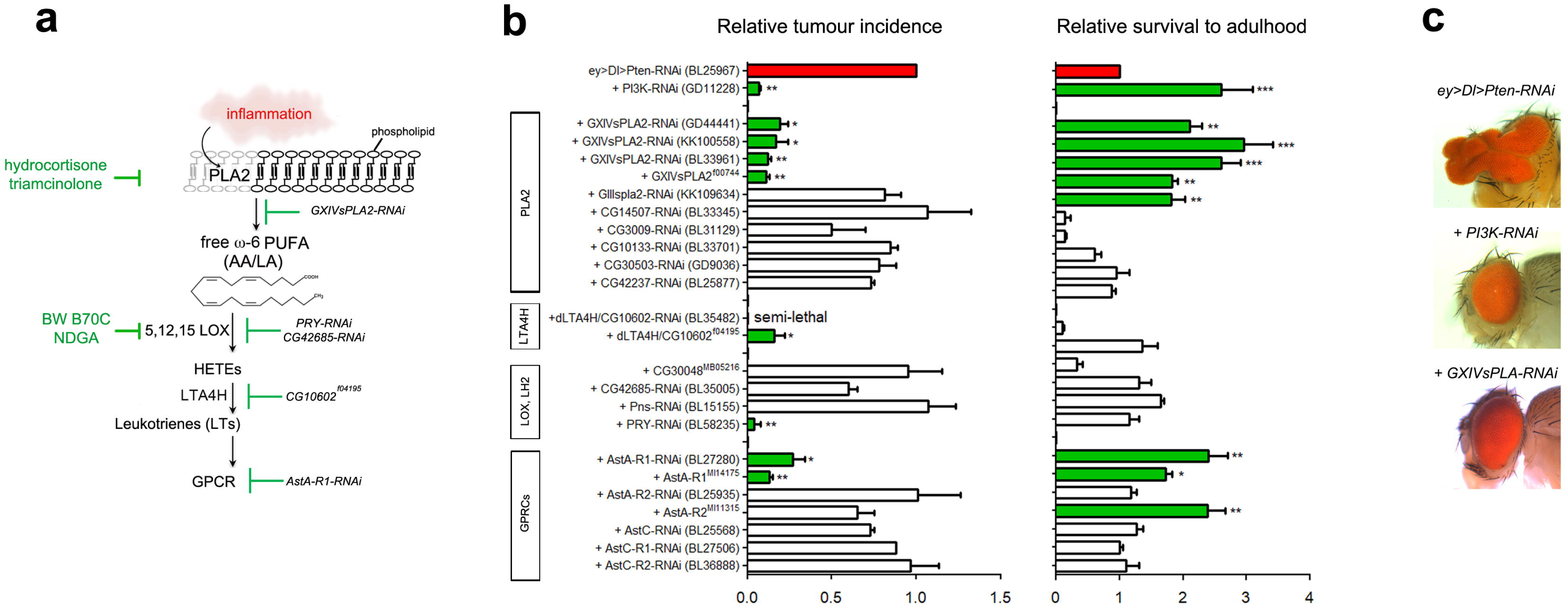
Genetic targeting of LOX signaling blocks Notch-PI3K/Akt cooperative oncogenesis. (**a**) Schematic LOX signalling pathway. Anti-tumorigenic drugs identified in our screen and RNAi-based silenced genes are shown. Homologous *Drosophila* genes (black) are also shown. In response to an inflammatory stimuli, PLA2 releases arachidonic acid and related ω-6 PUFAs from the membrane phospholipids. (**b**) Tumour rescue (left) and survival to adulthood (right) associated with *ey>Dl>Pten-RNAi*^*BL25967*^ after depleting the indicated genes via RNAi or mutation. Values (mean ± SD) are normalized with respect to *ey>Dl>Pten-RNAi*^*BL25967*^ (blue bar), and *>PI3K92E-RNAi*^*GD11228*^ acts as an internal positive control (c) Representative *ey>Dl>Pten-RNAi* plus RNAi for *PI3K92E* and *GXIVsPLA*. *n* = 50-100 eyes scored *per* genotype. \**P<*0.05, \*\**P<*0.01, \*\*\**P<*0.001, analysed by One-way ANOVA followed by Bonferroni’s Multiple Comparison test.

The most upstream step in the LOX-mediated production of pro-inflammatory lipid metabolites is the release of arachidonic acid mediated by the phospholipase A2 (PLA2) (Fig. 3a)^46^. Importantly, five suppressor drugs identified in our screen, including triamcinolone and hydrocortisone 21-hemisuccinate sodium, target this step (Supplementary Table 1 and Fig. 3a). In *Drosophila*, there are seven predicted PLA2 genes, all of which belong to the secretory class^47^. Tumour-specific RNAi silencing of *GXIVsPLA2* (a Group XII secretory PLA2), as well as halving its gene dosage (*GXIVsPLA2*^*f00744*^*/+*) suppressed tumorigenesis (Fig. 3b,c), essentially mirroring the anti-tumour effect of the drugs identified. Collectively, these data validate the screen results and uncovers a requirement of LOX-derivatives in Notch-PI3K/Akt-driven tumours.

### A pro-tumorigenic immune inflammation underlies Notch-PI3K/Akt cooperation

The discovery of a direct participation of NOS and LOX pathways in tumorigenesis promoted by Notch-PI3K/Akt interaction suggest an unanticipated connection between inflammation and this oncogenic cooperation. Inflammation has long been recognized as a contributing factor to solid cancer^48^, and can even be held responsible for treatment failure. Nevertheless, this strong correlation has not been addressed or observed in Notch-PI3K/Akt-driven tumorigenesis.

One of the main hallmarks of inflammation is macrophage infiltration^49^, and immune cells that infiltrate tumours can get involved in an active crosstalk with cancer cells^50^. Thus, we investigated whether macrophages are being recruited to the tumour site. Indeed, by using *GstD1-GFP* (an oxidative stress reporter expressed in haemocytes/macrophages) together with the haemocyte-specific marker *hmlΔ-DsRed*^51^ (Supplementary Fig. 4a), we confirmed the presence of infiltrating macrophages within tumorigenic tissue. We found that tissue resident macrophages showing a typical rounded morphology in wild type tissue (Supplementary Fig. 4b) became polarized (spindle shaped) in tumour tissues (Supplementary Fig. 4c), distinctive of activated macrophages. Interestingly, these tumour-associated macrophages (TAMs) are absent in BW B70C treated eye discs (Supplementary Fig. 4d,e). This data suggest that NOS/LOX activity shapes inflammatory macrophages response in Notch-PI3K/Akt tumours and links inflammation to tumorigenesis driven by these oncogenes.

A salient feature of pro-tumorigenic inflammation mediated by immune cells is immunosuppression^48,52,53^. In *Drosophila*, melanisation is a critical innate immune responses in the defence against tumour cells^54^, a process mediated by the prophenoloxidase (PPO)-activating cascade^55^. Therefore, to investigate the participation of inflammation-mediated immunosuppression in Notch-PI3K/Akt tumorigenesis we examined the expression of the *PPO* genes. While single *Notch* overexpression (*ey>Dl* hyperplastic eye discs) robustly stimulated *PPO1* and *PPO2* expression (Fig. 4a), both tumour-bearing larvae (*ey>Dl>Pten-RNAi*) and larvae with solely the PI3K/Akt pathway activated (*ey>Pten-RNAi*) prevented this response (Fig. 4a) indicating that activated PI3K/Akt signalling dampen the activity of the immune system. To get further insight on how immunosuppression modulates tumorigenesis, we genetically diminished PPO genes activity in *ey>Dl* larvae using a triple *PPO1-3* knockout^56^. Remarkable, we found that about 30% of emerging *ey>Dl*, *PPO1-3*^*–*^*/+* adults displayed tumorous eyes (Fig. 4b), similar to the results obtained by Notch-Nos over-expression (*ey>Dl>Nos,* Fig. 2d-e). These data indicate that immunosuppression is enough to release Notch oncogenic potential, a process prompted by the cooperation with activated PI3K/Akt.

**Figure 4.**
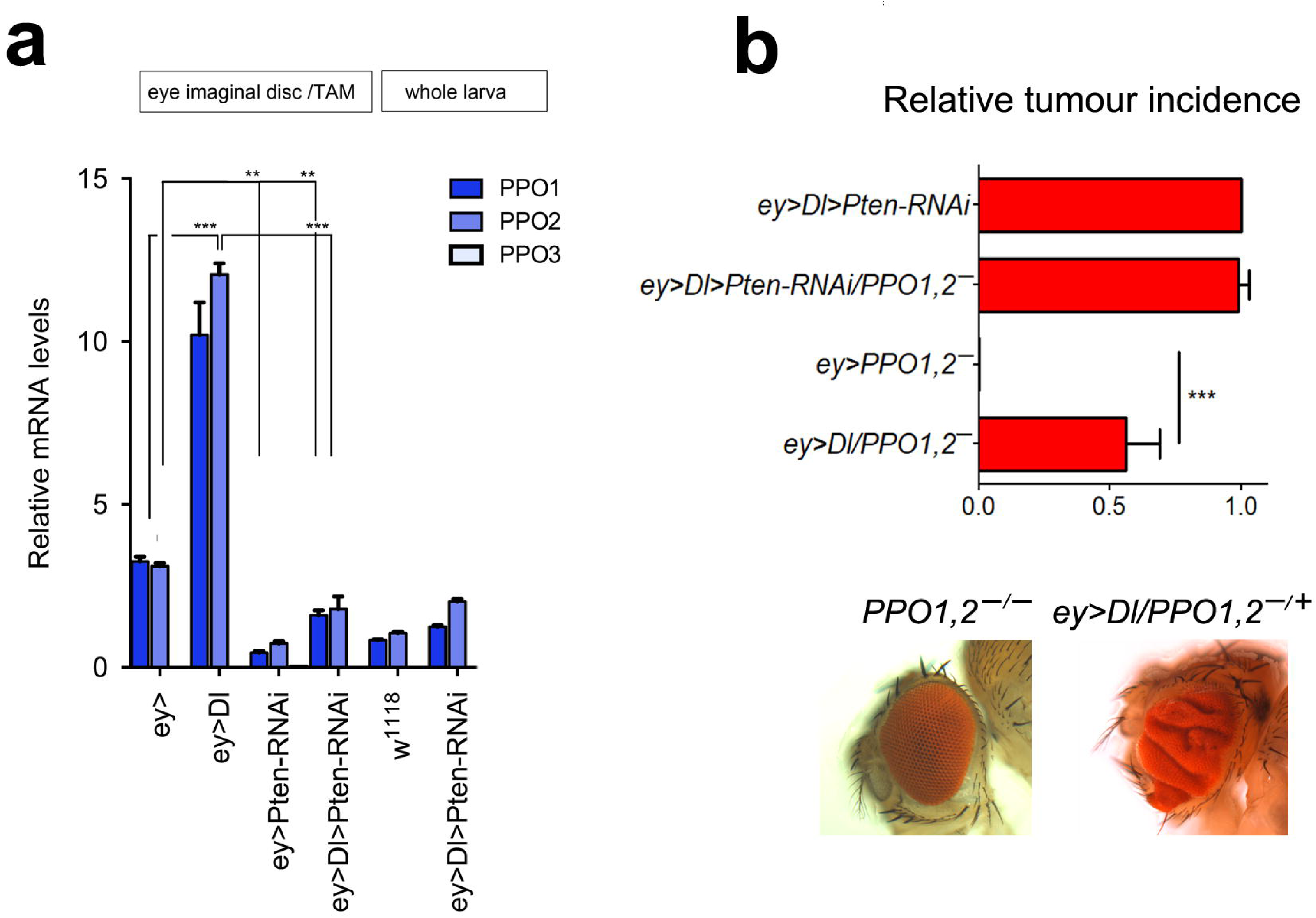
Immunosuppression. (**a**) Expression of the *PPO* genes in immune cells from larvae of the indicated genotypes analysed by qRT-PCR. Expression is shown for total larva (*n* = 5 per genotype) or for dissected eye imaginal discs (*n* = 30 eye discs per genotype). Experiments were performed in triplicate and data are presented as means ± SD. *PPO3* was undetected in these assays. Statistical significance was determined using a Student’s *t*-test. \**P<*0.05, ** *P<*0.01, *** *P<*0.001. (**b**) Relative tumour incidence (in %) in *ey>Dl;PPO1,2* (up) and representative eye phenotypes (down) (*n =* 50-100 eyes scored; \**P<*0.05, ** *P<*0.01, *** *P<*0.001). One-way ANOVA test, using Bonferroni multiple comparison tests.

## Discussion

How Notch and PI3K/Akt cooperate to perturb the physiological processes that regulate tissue growth, proliferation and metabolism to promote tumour initiation and progression remains poorly understood. Notch and PI3K/Akt inhibitors has been tested in several clinical trials and, although their use holds promise, their progress toward therapeutic treatment is hindered due to the side effects associated and drug resistance development^57-59^. Thus, therapies based on blocking directly Notch or PI3K/Akt activity needs to be thoroughly reconsidered. In this scenario, the characterization of targets and mechanism/s downstream of Notch and PI3K/Akt synergy is key to the identification of cancer vulnerabilities that can be therapeutically exploited.

Here, we report an *in vivo* drug discovery strategy in *Drosophila* that has led to the identification of pharmacologically active compounds with potential therapeutic impact. In particular, we found that BW B70C elicits a potent anti-tumorigenic response by blocking a NOS/LOX-dependent inflammatory mechanism. There are increasing evidence that points to a role for NOS^60,61^ and LOX^62-65^ in specific human oncogenic processes. Our data indicates that they are key pro-tumorigenic players underlying the synergy between Notch and PI3K/Akt signalling. Tumours can trigger an intrinsic inflammatory response that creates a pro-tumorigenic microenvironment^66^ that can play critical roles at different stages of cancer development^50^. We show here that diminishing the immune response or triggering an inflammatory program modulated by PI3K/Akt-dependent NOS signalling might be one mechanism to unleash Notch’s oncogenic potential. Moreover, the fact that NOS inhibition resulted in no harm for normal cells indicates that it can represent a promising safe druggable target for human cancers.

Finally, this study further highlights the considerable value that unbiased chemical screens in *Drosophila* have when it comes to deciphering new targets and potential novel therapeutic approaches in cancer.

## Acknowledgement

We thank B. Lemaitre, K. Brueckner and D. Bohmann for mutant flies and D. Ferres-Marco for the analysis of TSA in the ‘eyeful’ flies. We thank E. Ballesta-Illán, I. Oliveira, MC. Martinez-Moratalla, L. Mira, Csendesné Anna Rehák, Szilvia Bozsó and Anikó Berente for excellent technical assistance. We also thank the Bloomington Stock Center (NIH P40OD018537), the Drosophila Genomics Resource Center (NIH OD010949-10), the TRiP at Harvard Medical School (NIH/NIGMS R01-GM084947) for providing fly stocks. This work was supported by an FPI fellowship (BES-2015-073796) to LGL; the Hungarian Brain Research Program (KTIA_NAP_13-2-2014-0007) and the Hungarian Scientific Research Foundation (OTKA grant 109330) to JM; and the Generalitat Valenciana Grant (PROMETEO II/2013/001), the European Commission Grant (FP7-HEALH-F2-2008-201666), Foundation Botin (FB), and the Ministry of Economy and Competitiveness Grants (SEV-2013-0317 and MICCIN BFU2015-64239-R) to MD.

## Supplementary Figure legends

**Supplementary Figure 1. Proof-of-concept using Notch and PI3K inhibitors and formula to calculate drug response**. (**a**) Graphs show the tumour incidence of each fly models of Notch-PI3K/Akt cooperative oncogenesis. (b) Proof of principle of efficacy versus toxicity of drugs directly targeting individual notch or Pi3K/Akt signalling. The γ-secretase inhibitor DAPT {N-[N-(3,5-difluorophenacetyl)-l-alanyl]-S-phenylglycine t-butyl ester} and the PI3K inhibitor LY294002 work effectively in flies (Bjedov *et al*, 2010; Danilov *et al*, 2013; Micchelli *et al*, 2003). The graphs show the efficacy (tumour incidence) and toxicity (survival to adulhood) of adults emerging from drug fed tumour-bearing larvae overexpressing *Delta* and *dAkt1* by *ey-Gal4* at increasing doses (*n=*100 flies per drug and concentration). The mortality rate of untreated animals co-overexpressing *Delta* and *Akt* (or *Pten-RNAi*) by *ey-Gal4* is 79% (i.e. survival rate of 21%). Low-dose of DAPT rescued partially this tumour-associated mortality without reducing tumour burden. PI3K inhibitors caused 100% larval lethality, while non-lethal doses failed to suppress tumorigenesis. Combination of DAPT and LY294002 led to synergistic toxicity, resulting in high lethality even in the low-dose DAPT groups. (**c**) Formula to calculate response (R). The eye tumour phenotype can be influenced by culturing conditions such as humidity, temperature, and variation in the fly food. As such, vehicle-control groups grown in parallel were used to normalize the response. (**d**) Summary of results after first round (R1) of screening. 37% of the compounds (474/1280) showed a suppressor/enhancer response equal or higher than 20% and were selected for re-screening.

**Supplementary Figure 2. In vivo RNAi-based validations of drug screen results.** (**a**) Scheme of the genetic crosses to test *UAS-RNAi* transgenes of candidate drug targets *in vivo* in larvae carrying *UAS-Dl* and *UAS-Pten-RNAi* and *ey-Gal4*. F1 offspring was scored for the tumour burden and animal mortality. (**b**) Left: Bar graph shows tumour incidence (left) and survival to adulthood (right) after RNAi-mediated depletion of the predicted molecular drug target in the *ey>Dl>Pten-RNAi*>gene X-RNAi animals. (**c**) Representative eye tumour rescued phenotype carrying the indicated *UAS-RNAi* of candidate targets. We used the orthologs search tools from FlyBase, which includes Compara, eggNOG, Inparanoid, OMA, Panther, Phylome, RoundUp, TreeFam ortholog prediction algorithms.

**Supplementary Figure 3. Nos aberrant expression is induced by activated Akt.** (**a**) *Nos*^*MI09718/+*^ (*Nos sensor-GFP*, green) in control *ey-Gal4* eye discs is restricted to the postmitotic retinal cells and the optic nerve. Eye tumour discs show aberrant *Nos* in the undifferentiated proliferative eye region where *ey-Gal4* is expressed. (b) Relative mRNA levels of *dNOS* to *rp49* of the indicated genotypes (*n* = 30 eye discs per genotype) and (c) treated or not with BW B70C. One-way ANOVA test, using Bonferroni multiple comparison tests.

**Supplementary Figure 4. Tissue and tumour-associated macrophages and response to BW B70C.** (a,a’) Tissue resident macrophages are double labelled by GstD1-GFP (green), a sensitive oxidative stress reporter and the specific haemocyte/macrophage marker hmlΔdsRed (red). Discs are counterstained with DAPI (blue). (b) Tissue resident macrophages in wild type eye disc (ey>) and (c) in neoplastic Notch-PI3K/Akt tumoural disc in which TAMs shows a dispersed migratory spindle shape (arrows). (d) Representative polarized migratory hmlΔdsRed macrophages in Notch-PI3K/Akt tumoural disc (e) Treatment with BW B70C (20μg/ml) suppresses tumour growth and avoid macrophages polarization. (d’,e’) Magnifications of d and e showing spindle-shaped macrophages extending cell protrusions (arrows in d’) and rounded macrophages in e’.

### Materials and Methods

#### Drosophila husbandry

The full list of RNAi transgenes used is in Table EV4. In addition, the following fly strains were used: *w*^*1118*^, *ey-Gal4*, *ey-Gal4 UAS-Dl/CyO twist-GFP, ey-Gal4 UAS-Dl/CyO tub-Gal80, GS(2)1D233C (dAkt1)/TM2 twist-GFP, ey.Gal4 UAS-Dl GS(2)88A8*^*lola*^ ^*pipsqueak*^ (abbreviated *ey>Dl>Lola-Psq:* the eye tumour cancer strain), *Pten-RNAi* (BL25967) and *UAS-pTl.Venus* (BL30899) *UAS-Nos* (BL56830 and BL56823) from Bloomington Stock Centre, and *PI3K92E-RNAi (GD11228, v38985)* from the Vienna Drosophila RNAi Center, *GXIVsPLA2*^*f00744*^, *CG10602*^*f04195*^, *Pns*^*EY05553*^, *AstA-R1*^*MI14175*^ (*y*^*1*^ *w*^*\**^; *Mi{MIC} AstA-R1*^*MI14175*^), *Nos*^*MI09718*^ (*y*^*1*^ *w*^*\**^; *Mi{MIC}Nos*^*MI09718*^), *PKG/dg2*^*MI02855*^ (*y*^*1*^ *w*^*\**^; *Mi{MIC}dg2*^*MI0285*^), *PPO*^*Δ1-2,3*^ (a gift from B. Lemaitre), *GstD1-GFP* (a gift from D. Bohmann), *hmlΔ-DsRed* (a gift from K. Brueckner), *tub-Gal80, UAS-Dl* and *ey-Gal4* are described in FlyBase.

Flies were reared in standard fly food at 25ºC (except when indicated) on a 12-hour light/dark cycle. Standard ‘Iberian’ fly food was made by mixing 15 l of water, 0.75 kg of wheat flour, 1 kg of brown sugar, 0.5 kg yeast, 0.17 kg agar, 130 ml of a 5% nipagin solution in ethanol, and 130 ml of propionic acid.

#### Immunostaining and imaging of larval and adult eyes and wings

For assessing tumour suppression/enhancement by treatment, batches of late third instar larvae imaginal eye-antennal discs of treated or vehicle-control groups were dissected in PBS and collected in ice-chilled PBS. The tissue was fixed in 4% PFA at RT for 20 min and then washed three times with PBT (PBS buffer and 0.3% Triton). The tissue was incubated with DAPI (Invitrogen) for 15 min at RT (0.3 μg/ml), and washed again three times with PBT and a final wash with PBS. Discs were staining with anti-GFP, anti-DsRed and counterstaining with DAPI, and mounted in Vectashield (Vector Labs), and images were obtained using a Leica TCS SP2 Confocal microscope.

For adult wing notches analysis, adult wings from female flies were dissected and mounted on slides in 80% glycerol in phosphate-buffered saline solution. For imaging of adult eyes, flies were fixed and kept in 70% ethanol until imaging. Images were captured on an optical microscope ZEISS Axiophot, using a MicroPublisher 5.0 camera (QImaging) and the QCapture software (QImaging). All pictures were taken using a 5X objective with 1.5X zoom. Each eye image is a composite of 15 to 25 images of the same sample focused at different heights of the specimen. The in-focus composites were generated using the software AutoMontage Essentials 5.0.

#### Quantitative real-time PCR

Primers and probes for real-time (RT)-PCR were obtained from Applied Biosystems. Comparative RT-PCRs were performed in triplicates, and relative expression was calculated using the comparative Ct method. Primers were designed using the Primer3 online tool (http://bioinfo.ut.ee/primer3-0.4.0/primer3/). Data are presented as mean ± standard deviation; statistical analyses were performed using two-tailed Student’s *t*-test.

q-RT-PCR primer sequences

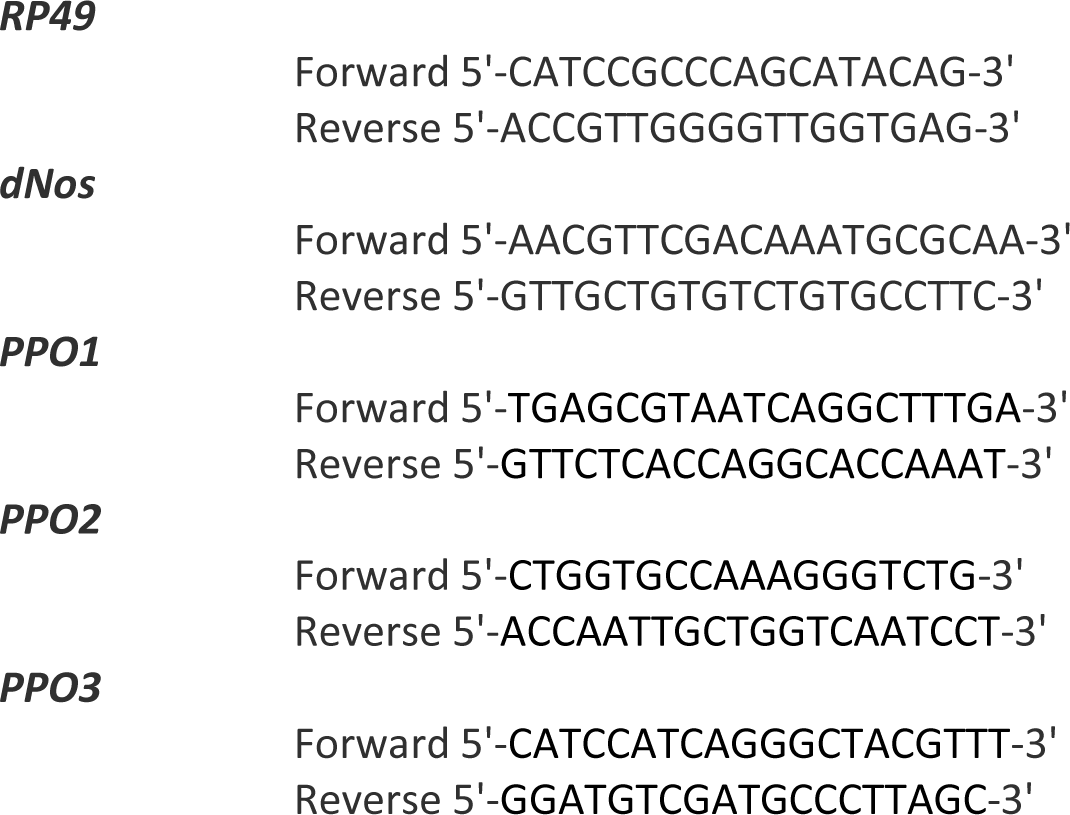

